# Atypical features of the disordered N-terminal region of a bacterial outer membrane copper transporter

**DOI:** 10.1101/2025.06.04.657848

**Authors:** Amira Khochtali, Marine Ote, Hugo Bâlon, Marc Dieu, Patricia Renard, Catherine Michaux, Jean-Yves Matroule

## Abstract

Metals like copper (Cu), zinc, and nickel exhibit dual nature, necessitating a tight regulation of their cellular homeostasis to meet physiological demands while preventing toxicity. In bacteria, metal homeostasis involves inner membrane (IM) P-type ATPases and ABC transporters, envelope-spanning tripartite efflux pumps, and outer membrane (OM) pore-forming proteins.

Four decades ago, the OM β-barrel protein PcoB was shown to provide an additional layer of Cu resistance in an *Escherichia coli* strain isolated from the gut of swine fed with Cu supplements. Interestingly, most PcoB homologs contain a poorly conserved disordered N-terminal domain (NTD) rich in histidine (His) and methionine (Met) residues, which are commonly associated with Cu coordination in cuproproteins. This suggests a potential role for the NTD in PcoB-mediated copper efflux.

We previously demonstrated that the free-living bacterium *Caulobacter vibroides* primarily relies on PcoB for Cu homeostasis. Here, we show that the NTD of *C. vibroides* PcoB is critical for PcoB function and stability, tolerating the swapping with the poorly conserved *E. coli* PcoB NTD and significant truncations. Unexpectedly, the predicted signal peptide (SP) was dispensable, challenging traditional concepts of protein translocation mechanisms. Moreover, the PcoB NTD plays a surprising role in stabilizing the periplasmic multicopper oxidase PcoA, encoded within the same operon as PcoB, highlighting a new role for an intrinsically disordered region (IDR).

**Importance:** Bacterial copper (Cu) homeostasis is essential for survival in fluctuating environments, yet the role of single outer membrane proteins in this process remains poorly characterized. Our study reveals that the intrinsically disordered N-terminal region of the β-barrel protein PcoB in *Caulobacter vibroides* plays a critical role in Cu efflux and PcoB stability. Remarkably, this region harbors an atypical signal peptide and contributes to the stability of the periplasmic multicopper oxidase PcoA, suggesting a novel regulatory function for an intrinsically disordered region. These findings challenge existing paradigms of protein targeting and homeostasis, with broad implications for understanding bacterial adaptation to stress.

## Introduction

Since the 19^th^ century, copper (Cu) has been widely recognized as an effective antimicrobial agent due to its toxicity at high concentration, which mainly results from mismetallation events (1), reactive oxygen species (ROS) production (2, 3) and protein aggregation (4).

Bacteria often encounter Cu in their natural environments, where it can be found in sediments or accumulate in mammalian macrophages as a defense mechanism against bacterial infections (5–7). Consequently, bacteria have evolved Cu resistance strategies to survive these harsh conditions (8). The most conserved Cu resistance strategies include (a) intracellular or extracellular Cu ions chelation by metallothioneins, bufferins and glutathione for instance (9, 10), (b) oxidation of Cu(I) ions to the less toxic Cu(II) ions by multicopper oxidases such as CueO (11), and (c) Cu efflux. In Gram-negative bacteria, Cu efflux is achieved through the combined action of (i) inner membrane (IM) P1B-type ATPases that transport Cu(I) ions from the cytoplasm to the periplasm (12), and (ii) tripartite HME RND pumps such as CusABC spanning the bacterial envelope and use the proton motive force (PMF) to expel periplasmic and potentially cytoplasmic Cu(I) out of the cell with the help of a periplasmic chaperone such as CusF(13).

In the early 1980s, an additional seven-component Cu resistance system was identified in an *Escherichia coli* strain isolated from the feces of swine fed with a Cu-rich diet (14, 15). This system, known as the Pco system, is encoded by a plasmid-born operon and has been poorly characterized so far. Among its components, only the multicopper oxidase PcoA has been identified as a close homolog of the CueO multicopper oxidase (16). The other Pco components are believed to be involved in periplasmic Cu chelation and efflux into the extracellular medium (16).

Recently, we have shown that the aquatic α-proteobacterium *Caulobacter vibroides* harbors a chromosomal two-gene operon encoding the PcoA multicopper oxidase and the outer membrane (OM) β-barrel protein PcoB, both involved in Cu resistance (17). PcoA and PcoB are expressed in sessile stalked cells and facilitate rapid Cu detoxification and efflux, favoring DNA replication and cell division (17).

Recently, we have experimentally and bioinformatically shown that PcoB preferentially binds Cu(II) ions and assembles into a β-barrel harboring a disordered N-terminal tail (18).

Given the OM location of PcoB, the mechanism underlying PcoB-mediated Cu efflux remains unclear and very likely differs from the well-known metal efflux ATPases and HME RNDs that are powered by ATP hydrolysis and the PMF, respectively.

In the present study, we investigated the role of the disordered N-terminal region of PcoB in Cu resistance. We provided evidence that this region contributes to PcoB’s function and stability and more surprisingly to PcoA stability, highlighting a new role for an intrinsically disordered region (IDR). In addition, PcoB seems to harbor an atypical signal peptide (SP), challenging existing paradigms on protein translocation across the IM.

## Results

### The disordered PcoB N-terminal domain is required for Cu resistance

The predicted 3D structure of PcoB by AlphaFold3 is consistent with its OM location and reveals two key features: (1) a C-terminal domain (CTD) organized into β sheets with 12 antiparallel β-strands forming a small β-barrel, and (2) a 109-residue-long IDR as the N-terminal domain (NTD) with one short α-helix encompassing residues 91 to 104 **(Fig. 1A**). This predicted structure aligns with the crystal structures of the PcoB β-barrel from *E. coli* and *Acinetobacter baumannii* which were resolved at a 2.0 Å and 6.5 Å resolution, respectively (19, 20). However, the high flexibility of the NTD prevented the crystallization of the full protein, which was therefore limited to the β-barrel.

**Figure 1.**
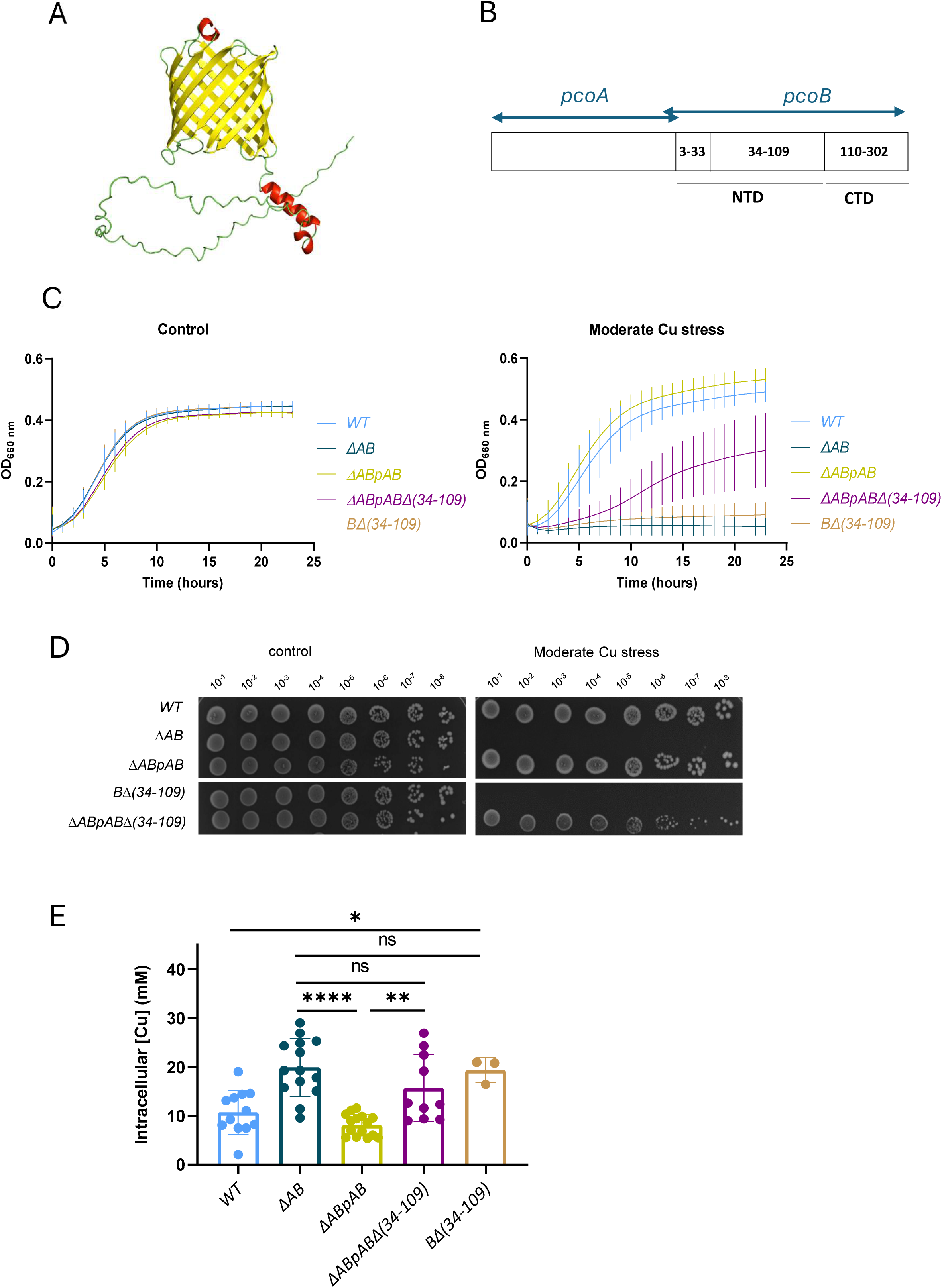
The disordered PcoB N-terminal domain is required for Cu resistance. **A.** Structure prediction of PcoB using AlphaFold3. **B.** Organization of *pcoAB* operon. **C.** Growth profiles at an absorbance of 660 nm of WT*, ΔAB, ΔABpAB, ΔABpABΔ(34-109)* and *BΔ(34-109)* strains grown in PYE medium (left) and in PYE medium supplemented with CuSO_4_ (right). Mean ± SD, at least three biological replicates. **D.** Viability assay on PYE plates of WT*, ΔAB, ΔABpAB, BΔ(34-109)* and *ΔABpABΔ(34-109)* strains, in control and moderate CuSO_4_ stress condition. **E**. Average Cu concentration in each bacterial cell exposed to 175 µM CuSO_4_ for 5 min. Mean ± SD, at least three biological replicates. *p* values were calculated using ANOVA combined with Dunnett’s multiple comparison test (\**p* < 0.05, ** *p* < 0.01, *** *p* < 0.001 and \*\*\*\**p* < 0.0001).

In *C. vibroides*, the PcoB intrinsically disordered NTD represents more than 30% of total protein sequence, classifying PcoB as a hybrid intrinsically disordered protein (IDP) (18). PcoB NTD also contains 17 His and 5 Met residues and is very likely oriented towards the periplasm (18). His and Met residues are frequently involved in Cu coordination (21), suggesting that the PcoB NTD may bind Cu and facilitate Cu efflux.

As an OM protein, PcoB is supposed to harbor a SP for translocation through the IM via the Sec translocon. Surprisingly, no SP could be detected in PcoB using the signal peptide prediction tool SignalP-5.0 **(Fig. S1A)**. Despite the extensive diversity of bacterial Sec signal peptides, certain features are conserved in all Sec substrates, including a three-region design: a positively charged N-terminal region (n-region), a hydrophobic central region (h-region), and a neutral, polar C-terminal region (c-region), along with a three-residue motif for signal peptidase I, typically comprising AXA motifs at the end of the c-region (22). Manual analysis of the PcoB sequence revealed a potential poorly conserved Sec motif, characterized by a high occurrence of hydrophobic residues and AXA patterns at positions 13-15, 20-22, and 34-36 **(Fig. S1B)**. To confirm this prediction, we isolated a periplasmic/OM fraction from exponentially growing *C. vibroides* cells expected to contain the mature PcoB and subjected this fraction to trypsin digestion followed by Liquid Chromatography/Mass Spectrometry (LC/MS) analysis. No peptide corresponding to the first 33 residues of PcoB was detected by LC/MS, suggesting that this peptide was cleaved during PcoB translocation through the IM via the Sec translocon **(Fig. S1C)**. Therefore, we decided to include the first 33 residues of PcoB as a putative SP in the subsequent genetic constructs to allow correct PcoB addressing to the OM. To investigate the role of the PcoB NTD_34-109_ in Cu resistance, we deleted the region encompassing the Alanine 34 to Threonine 109 residues within the chromosomal *pcoB* allele (*BΔ(34-109)*) or within the extrachromosomal *pcoAB* operon cloned in a low-copy pMR10 plasmid under the control of the strong and constitutive pLac promoter (*ΔABpABΔ(34-109)*) **(Fig. 1B)**. The growth of the resulting mutant strains was monitored over 24 h in liquid rich PYE medium under low (50 µM), moderate (100 µM CuSO_4_), and high (150 µM CuSO_4_) Cu stress conditions. The *ΔABpABΔ(34-109)* mutant exhibited an increased Cu sensitivity under moderate Cu stress relative to the WT and the complemented *ΔABpAB* genetic backgrounds **(Fig. 1C)**. This Cu sensitivity was concentration-dependent, with only a minor growth delay under low Cu stress and complete growth inhibition under high Cu stress **(Fig. S2A)**.

The increased Cu sensitivity of the *ΔABpABΔ(34-109)* mutant was also observed on plate, where diluted cultures were spotted on solid PYE medium under moderate (75 μM CuSO_4_) **(Fig. 1D)** and high (100 μM CuSO_4_) **(Fig. S2B)** Cu stress. The *BΔ(34-109)* chromosomal mutant displayed a ΔAB mutant phenotype even under low Cu stress, suggesting a higher impact of the *BΔ(34-109)* allele when present as a single chromosomal copy **(Fig. 1C-D)**.

To determine whether the increased Cu sensitivity of the *BΔ(34-109)* and *ΔABpABΔ(34-109)* mutants results from a defect in Cu efflux, we measured the Cu content in total cell extracts using ICP-OES.

Consistent with the observed Cu sensitivity, the *BΔ(34-109)* and the *ΔABpABΔ(34-109)* mutants displayed a significant increase of the intracellular Cu levels compared to the WT and *ΔABpAB* control strains, respectively, and comparable to the *ΔAB* mutant **(Fig. 1E)**, indicating that the PcoB NTD_34-109_ is crucial for Cu efflux.

### PcoB and PcoA stability relies on the PcoB NTD

The increased sensitivity of the *BΔ(34-109)* and *ΔABpABΔ(34-109)* mutants could result from a loss-of-function and/or reduced stability of the mutated PcoB. Consistent with this latter hypothesis, PcoB remained undetectable when performing an anti-PcoB immuno-blot on total cell extracts from the *BΔ(34-109)* and the *ΔABpABΔ(34-109*) mutants **(Fig. 2A)**.

**Figure 2.**
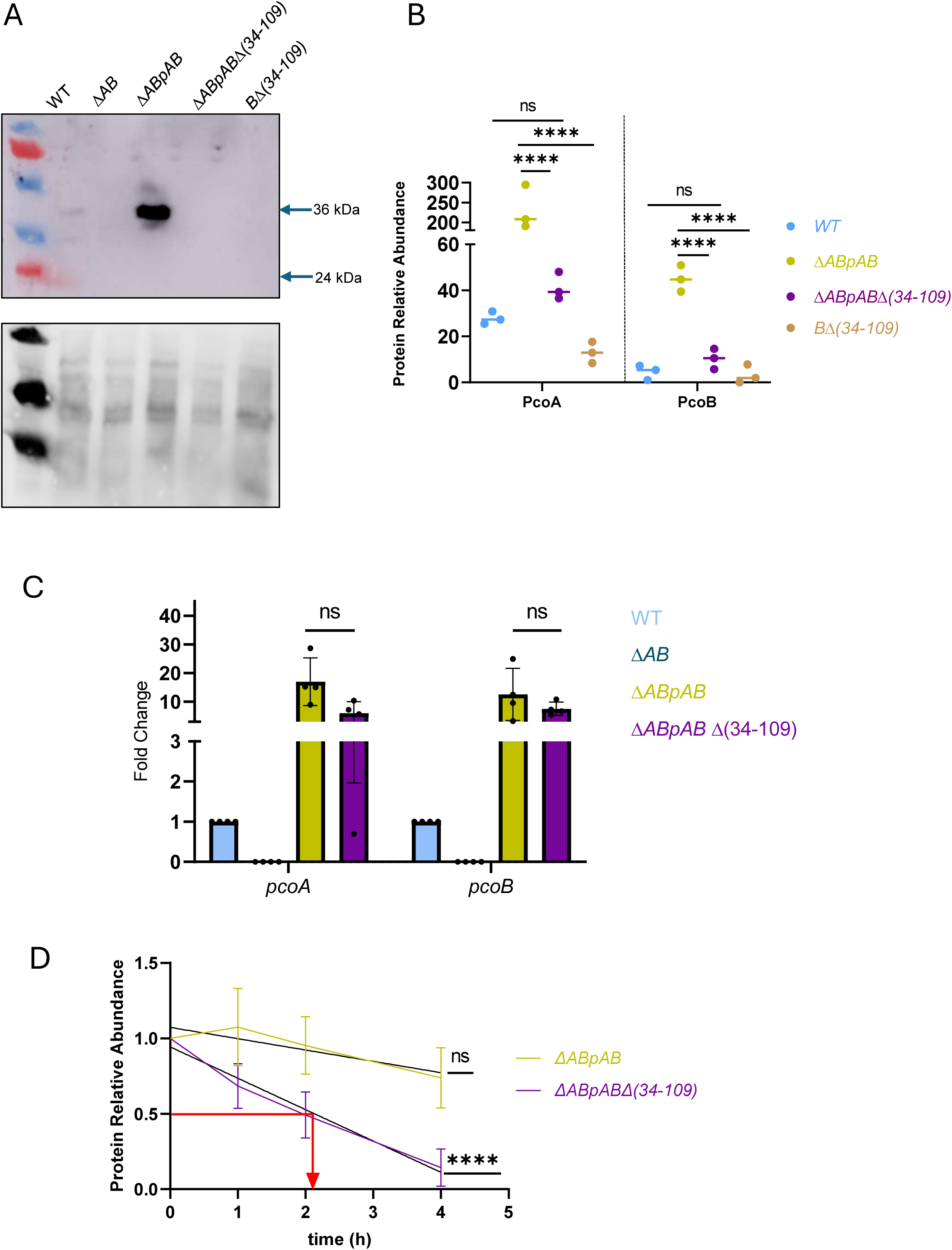
PcoB and PcoA stability relies on the PcoB NTD. **A.** Immunoblot anti-PcoB performed on total cell extracts from WT*, ΔAB, ΔABpAB, ΔABpABΔ(34-109)* and *BΔ(34-109)* strains grown in PYE medium. The lower panel corresponds to the Cy5-stained gel used as a loading control. **B**. Normalized spectrum counts of PcoA and PcoB peptides in WT*, ΔABpAB, ΔABpABΔ(34-109)* and *BΔ(34-109)* strains grown in PYE medium, measured by LC-MS. Individual values and means represented. **C.** Relative *pcoA* and *pcoB* mRNA level measured by RT-qPCR in the WT*, ΔAB, ΔABpAB, and ΔABpABΔ(34-109)* strains grown in PYE medium. *p* values were calculated using ANOVA combined with Dunnett’s multiple comparison test (\**p* < 0.05, ** *p* < 0.01, *** *p* < 0.001 and \*\*\*\**p* < 0.0001). **D**. PcoA abundance measured by LC/MS in total cell extracts obtained 0 h, 1 h, 2 h and 4 h after chloramphenicol treatment of the *ΔABpAB and ΔABpABΔ(34-109)* strains grown in PYE medium. *p* values were calculated using a linear regression statistical test; \*\*\*\**p* < 0.0001.

To rule out any epitope loss in the PcoBΔ(34-109) mutant that could impair anti-PcoB antibody recognition and immuno-blot reliability, we quantified PcoB in periplasmic/OM fractions using LC/MS. Consistent with the immuno-blot data, the deletion of the PcoB NTD (34-109) domain leads to a 4.4-fold decrease of PcoB relative to the *ΔABpAB* strain used as a control **(Fig. 2B)**. Yet, PcoB abundance remained at a WT level in the *ΔABpABΔ(34-109)* mutant. Considering the low expression of the chromosomal *pcoB*, we decided to mainly focus on the plasmidic version of PcoB, where *pcoB* was kept in operon with *pcoA* to ensure the coupled translation of PcoA and PcoB. Indeed, in the *ΔABpB* strain where the sole *pcoB* gene is present on pMR10, PcoB abundance is lower than in the *ΔABpAB* strain (**Fig. S3**).

Unexpectedly, the LC/MS analysis also revealed a significant 18-fold and 5.6-fold decrease in PcoA levels in the *BΔ(34-109)* and in the *ΔABpABΔ(34-109)*, respectively **(Fig. 2B)**. Accordingly, PcoA abundance is also reduced in the *ΔABpA* background (relative to the *ΔABpAB* background), which mimics a *ΔB* mutant (**Fig. S3**). This latter observation suggests that PcoB, and more specifically the PcoB NTD_34-109_ is required to sustain PcoA homeostasis.

It is unlikely that the decrease in PcoA and PcoB abundance observed in the *ΔABpABΔ(34-109)* mutant results from a down-regulation of the *pcoAB* operon transcription, considering that the *pcoAB* operon is under the control of the constitutive lac promoter when expressed from the pMR10 plasmid. Consistent with this assumption, the relative amounts of *pcoA* and *pcoB* transcripts measured by RT-qPCR did not significantly vary between the *ΔABpABΔ(34-109)* mutant and the *ΔABpAB* strain **(Fig. 2C)**.

To test whether the PcoB NTD_34-109_ could play a role in PcoA and PcoB stability, we aimed to compare their respective half-life in the *ΔABpABΔ(34-109)* and *ΔABpAB* backgrounds. PcoA and PcoB levels were measured by LC/MS in total protein extracts at different timepoints after translation inhibition with 100 μg/ml chloramphenicol. In the *ΔABpAB* strain, PcoA is relatively stable, with protein levels remaining constant over 4 h after translation inhibition **(Fig. 2D)**. However, in the *ΔABpABΔ(34-109)* mutant, PcoA content rapidly decreases after protein translation arrest to a level of 50% after 2 h, indicating that the PcoB NTD_34-109_ contributes to PcoA stability **(Fig. 2D)**.

Interestingly PcoB stability was hardly affected by the deletion of the PcoB NTD_34-109_, at least during the first 4 hours of protein translation arrest, suggesting that the lower abundance of the PcoBΔ(34-109) does not result from a post-translational destabilization as it is observed for PcoA **(Fig. S4)**.

Considering that PcoA and PcoB protein levels in the *ΔABpABΔ(34-109)* mutant remain equivalent to the levels measured in the WT strain **(Fig. 2B)**, we propose that the Cu sensitivity of the *ΔABpABΔ(34-109)* mutant is, at least partially, due to a complete or partial loss of PcoB function.

### PcoB NTD (34-109) domain is tolerant to sequence and size change

The alignment of the *C. vibroides* PcoB protein sequence with PcoB orthologs from α, β and γ-proteobacteria reveals a poor conservation of PcoB NTD compared to the well-conserved CTD forming the β-barrel **(Fig. 3A)**. However, the disorder of PcoB NTD seems to be present in all PcoB orthologs **(Fig. 3B-C)**, suggesting that the role of the PcoB NTD may not be directly related to its specific amino acid composition but rather to its length and/or disordered structure.

**Figure 3.**
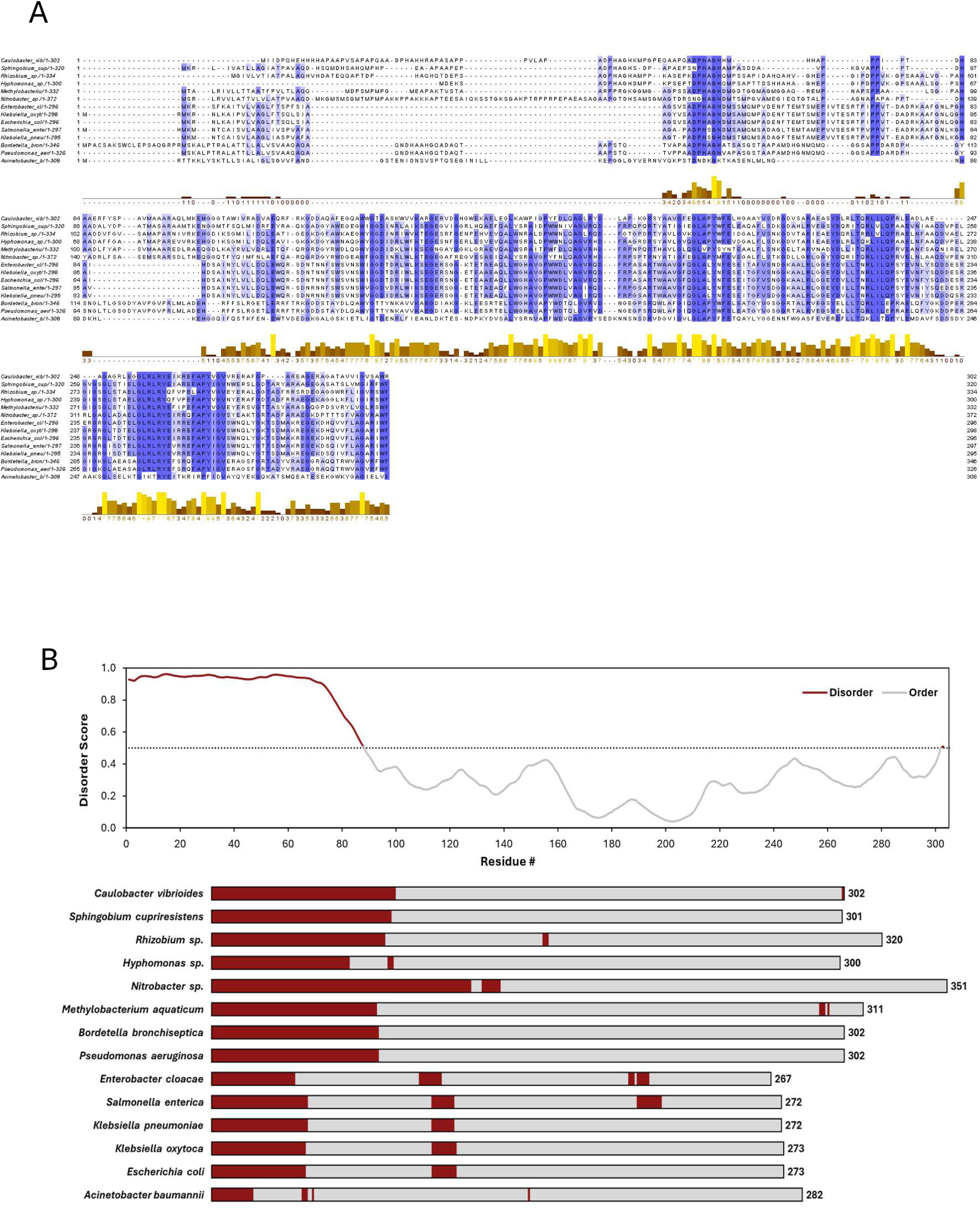
PcoB NTD is poorly conserved. **A.** Multiple MAFFT alignment of PcoB_Cc_ with 13 PcoB homologs from α-, β-, δ- and γ-proteobacteria. Identical residues are highlighted in blue. Predicted conservation and intrinsic disorder profile of PcoB homologs. **B.** RIDAO mean disorder profile (MDP) of *Caulobacter vibrioides* PcoB. Every residue displaying a score above the 0.5 threshold is considered disordered. **C.** Schematic representation of MDP derived from RIDAO for 13 homologs of *C. vibrioides* PcoB identified by sequence alignment and conservation. Disordered and ordered regions are highlighted in dark red and grey, respectively.

To test this hypothesis, we swapped the NTD_34-109_ of *C. vibroides* PcoB with the NTD_25-90_ of *E. coli* PcoB (*ΔABpABN_Ec_*), keeping the first 33 residues from *C. vibroides* PcoB as a SP, **(Fig. 4A)** and tested the ability of the resulting chimeric PcoB to sustain Cu resistance. The *ΔABpABN_Ec_* strain shows similar growth pattern to the *ΔABpAB* strain under control conditions and under any of the tested Cu stress conditions in liquid medium **(Fig. 4B)** or on plates **(Fig. 4C)**. Consistent with this observation, the *ΔABpABN_Ec_* mutant displayed an intracellular Cu content equivalent to that of the *ΔABpAB* strain and significantly lower than the *ΔAB* mutant **(Fig. 4D)**. Accordingly, the PcoB protein levels were not affected by the domain swapping, whereas PcoA levels were significantly reduced, yet remaining much higher than in the *ΔABpABΔ(34-109)* mutant (Fig. 4E).

**Figure 4.**
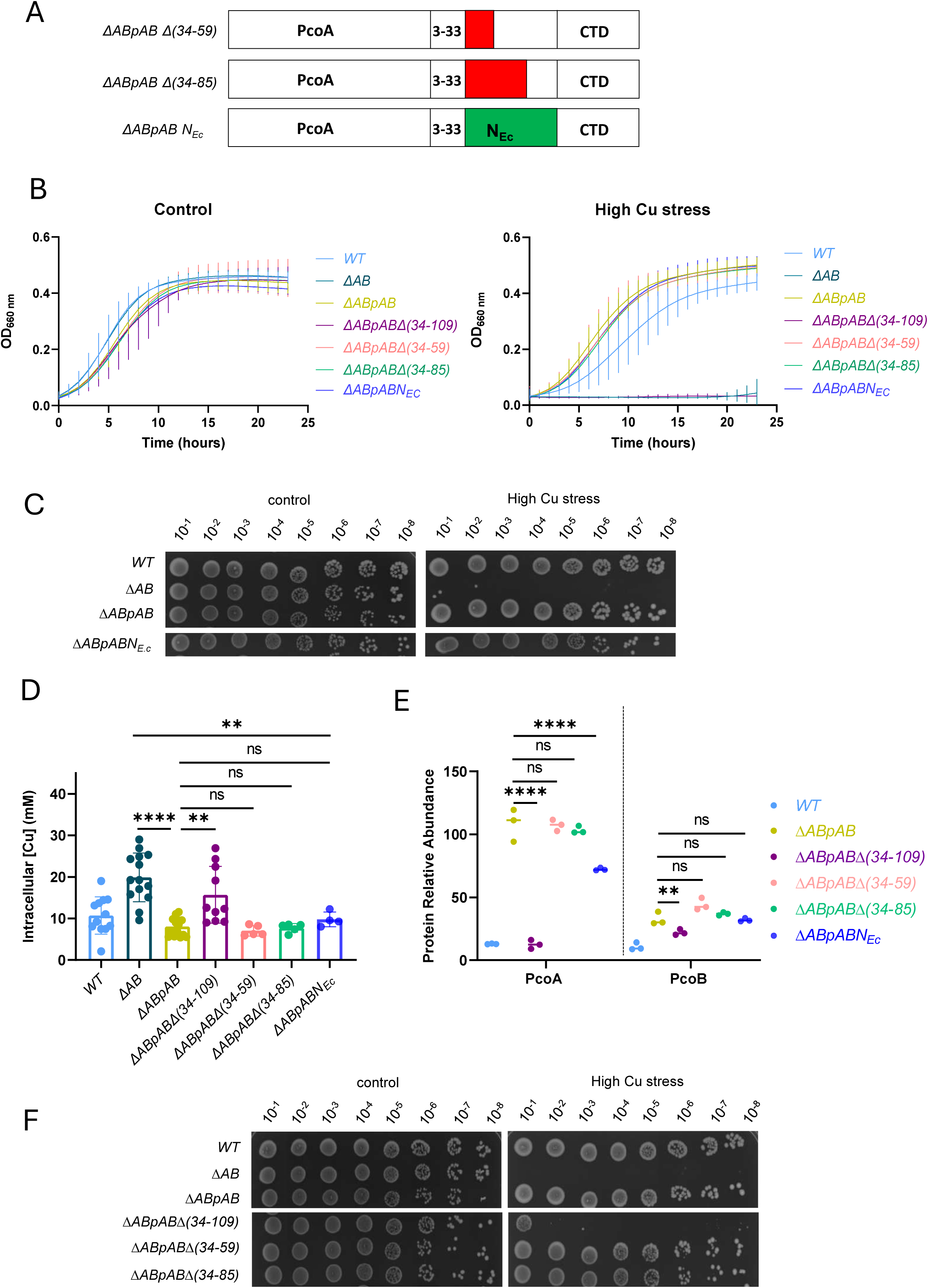
PcoB NTD is tolerant to sequence and size change. **A.** schematic representation of the *ΔABpABΔ(34-59)*, *ΔABpABΔ(34-85)* and *ΔABpABN_Ec_* mutants highlighting the deleted regions (in red) or the swapped region (green). **B.** Growth profiles at an absorbance of 660 nm of WT*, ΔAB, ΔABpAB, ΔABpABΔ(34-109), ΔABpABΔ(34-59), ΔABpABΔ(34-85)* and *ΔABpABN_Ec_* strains grown in PYE medium (left) and in PYE medium supplemented with CuSO_4_ (right). Mean ± SD, at least three biological replicates. **C.** Viability assay on PYE plates of WT*, ΔAB, ΔABpAB and ΔABpABN_Ec_* strains, in control and high CuSO_4_ stress conditions. **D**. Average Cu concentration in each bacterial cell in the WT*, ΔAB, ΔABpAB, ΔABpABΔ(34-109), ΔABpABΔ(34-59), ΔABpABΔ(34-85)* and *ΔABpABN_Ec_* strains exposed to 175 µM CuSO_4_ for 5 min. Mean ± SD, at least three biological replicates. **E.** Normalized spectrum counts of PcoA and PcoB peptides in the WT*, ΔABpAB, ΔABpABΔ(34-109), ΔABpABΔ(34-59), ΔABpABΔ(34-85), and ΔABpABN_Ec_* strains grown in PYE medium, measured by LC-MS. Individual values and means represented. *p* values were calculated using ANOVA combined with Dunnett’s multiple comparison test (\**p* < 0.05, ** *p* < 0.01, *** *p* < 0.001 and \*\*\*\**p* < 0.0001). **F.** Viability assay on PYE plates of WT*, ΔAB, ΔABpAB, ΔABpABΔ(34-109), ΔABpABΔ(34-59) and ΔABpABΔ(34-85)* strains, in control and high CuSO_4_ stress conditions.

Together, these results indicate that the overall primary sequence of the PcoB NTD_34-109_ does not seem to be essential for PcoB stability and function.

Considering the apparent tolerance of the PcoB NTD_34-109_ to sequence change, we questioned whether the length of this disordered N-term domain might be a limiting factor. To address this point, we generated the *ΔABpABΔ(34-59)* and *ΔABpABΔ(34-85)* mutants, missing the first 26 and 52 residues, respectively **(Fig. 4A)**.

Surprisingly, these mutants phenocopied the *ΔABpA*B strain in liquid medium **(Fig. 4B)** and on plates **(Fig. 4F)** under any of the tested Cu stress conditions. Consistent with these observations, the intracellular Cu content **(Fig. 4D)** and the PcoA and PcoB levels **(Fig. 4E)** of these mutants was comparable to the *ΔABpAB* strain under Cu stress. These results indicate that the PcoB NTD_34-109_ can be significantly shortened without affecting the stability of PcoB and PcoA or the function of PcoB.

### Cu resistance relies on a minimal set of His residues within PcoB NTD

The presence of 9 His and 4 Met residues in the PcoB NTD_34-109_ **(Fig. 5A)** led us to hypothesize that these residues may bind Cu ions and play a key role in PcoB function. Therefore, we mutated these His (H) or Met (M) residues into Ala (A), yielding the *ΔABpABH49-105A* (H49, H52, H67, H70, H71, H73, H74, H83, and H105) and the *ΔABpABM54-102A* (M54, M72, M94, and M102) mutants **(Fig. 5B)**.

**Figure 5.**
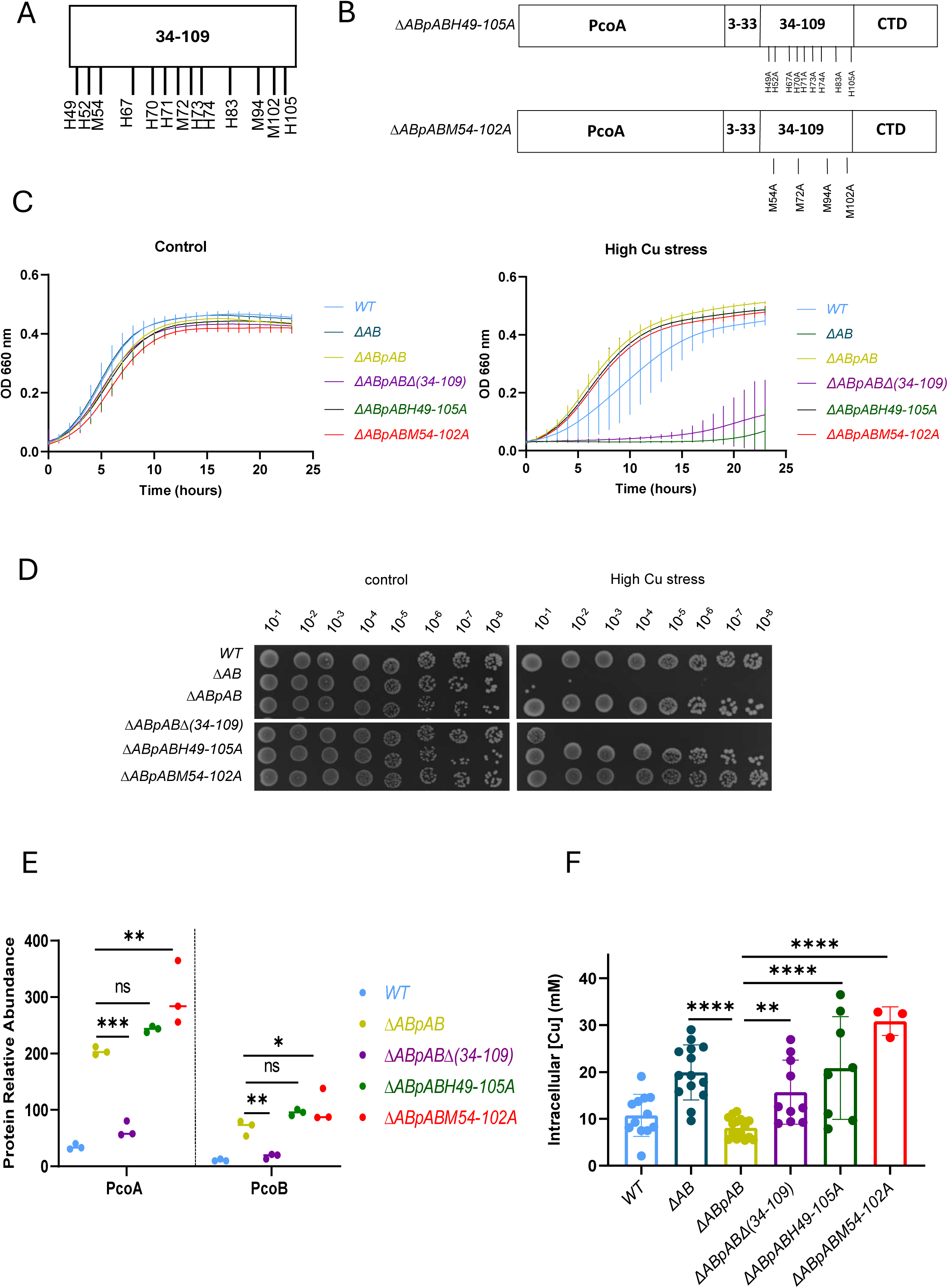
Met and His residues of PcoB NTD play a key role in PcoB function. **A.** schematic representation of the His and Met residues located in the PcoB 34-109region. **B.** schematic representation of the *ΔABpABH49-105A* and *ΔABpAM54-102A* mutants. **C.** Growth profiles at an absorbance of 660 nm of WT*, ΔAB, ΔABpAB, ΔABpABΔ(34-109), ΔABpABH49-105A* and *ΔABpABM54-102A* strains grown in PYE medium (left) and in PYE medium supplemented with CuSO_4_ (right). **D.** Viability assay on PYE plates of WT*, ΔAB, ΔABpAB, ΔABpABΔ(34-109)*, *ΔABpABH49-105A* and *ΔABpABM54-102A* strains, in control and high CuSO_4_ stress conditions. **E.** Normalized spectrum counts of PcoA and PcoB peptides in the WT*, ΔABpAB, ΔABpABΔ(34-109), ΔABpABH49-105A* and *ΔABpABM54-102A* strains grown in PYE medium, measured by LC-MS. Individual values and means represented. **F**. Average Cu concentration in each bacterial cell in the WT*, ΔAB, ΔABpAB, ΔABpABΔ(34-109)*, *ΔABpABH49-105A* and *ΔABpABM54-102A* strains exposed to 175 µM CuSO_4_ for 5 min. Mean ± SD, at least three biological replicates. *p* values were calculated using ANOVA combined with Dunnett’s multiple comparison test (\**p* < 0.05, ** *p* < 0.01, *** *p* < 0.001 and \*\*\*\**p* < 0.0001)

These multiple point mutants exhibited a similar growth profile to the *ΔABpAB* strain under both control and Cu stress conditions **(Fig. 5C)**, which was further confirmed on plate **(Fig. 5D)**. Consistent with these observations, the PcoA and PcoB level were not negatively affected by the point mutations **(Fig. 5E)**. Surprisingly, mutating the His and the Met residues in the PcoB NTD_34-109_ led to a striking Cu accumulation, suggesting that these residues play a key role in PcoB-dependent Cu efflux **(Fig. 5F)**. The lack of Cu sensitivity of these multiple point mutants might therefore be due to the high level of PcoA detoxifying the high amount of accumulated Cu.

The presence of 8 His residues within the predicted SP is puzzling and not observed in the other PcoB orthologs **(Fig. 3A).** To address this peculiarity, we constructed the *ΔABpABΔ(3-33)* mutant, lacking the putative SP sequence **(Fig. 6A)**. This *ΔABpABΔ(3-33)* mutant displays a *ΔABpAB*-like Cu resistance in both liquid medium **(Fig. 6B)** and on plate **(Fig. 6C)**. Furthermore, this mutation does not impact Cu content **(Fig. 6D)** and PcoA level **(Fig. 6E)** but reduces PcoB level in the periplasmic/OM fraction, even if it remains at a much higher level than in the WT strain **(Fig. 6E)**. PcoB levels measured in purified OM fraction from the *ΔABpABΔ(3-33)* mutant and the control *ΔABpAB* strains indicate that the PcoB NTD_3-33_ is not required to address PcoB at the OM **(Fig. 6F)** and prompted us to question its role as a canonical SP. To test this hypothesis, we repeated the LC/MS analysis on a periplasmic/OM fraction, combining trypsin and GluC peptidase, the latter cleaving after glutamate (E) and aspartate (D) residues, potentially generating peptides within the PcoB (3–33) region. Indeed, the trypsin peptidase (used to digest the proteins before the LC/MS) cleaves after Arginine (R) and lysine (K), and the first cleavage site within PcoB aligns with the end of the putative SP (R_33_), which is not optimal for generating peptides within the SP sequence for detection and sequencing. This adapted digestion revealed the presence of the 9-33 peptide in the mature form of PcoB **(Fig. S5)**, reinforcing the idea that PcoB does not harbor a conventional SP in *C. vibroides*, which contrasts with most of the PcoB homologs analyzed in this study, but *Hyphomonas* sp., predicted to harbor a conventional SP by SignalP.

**Figure 6.**
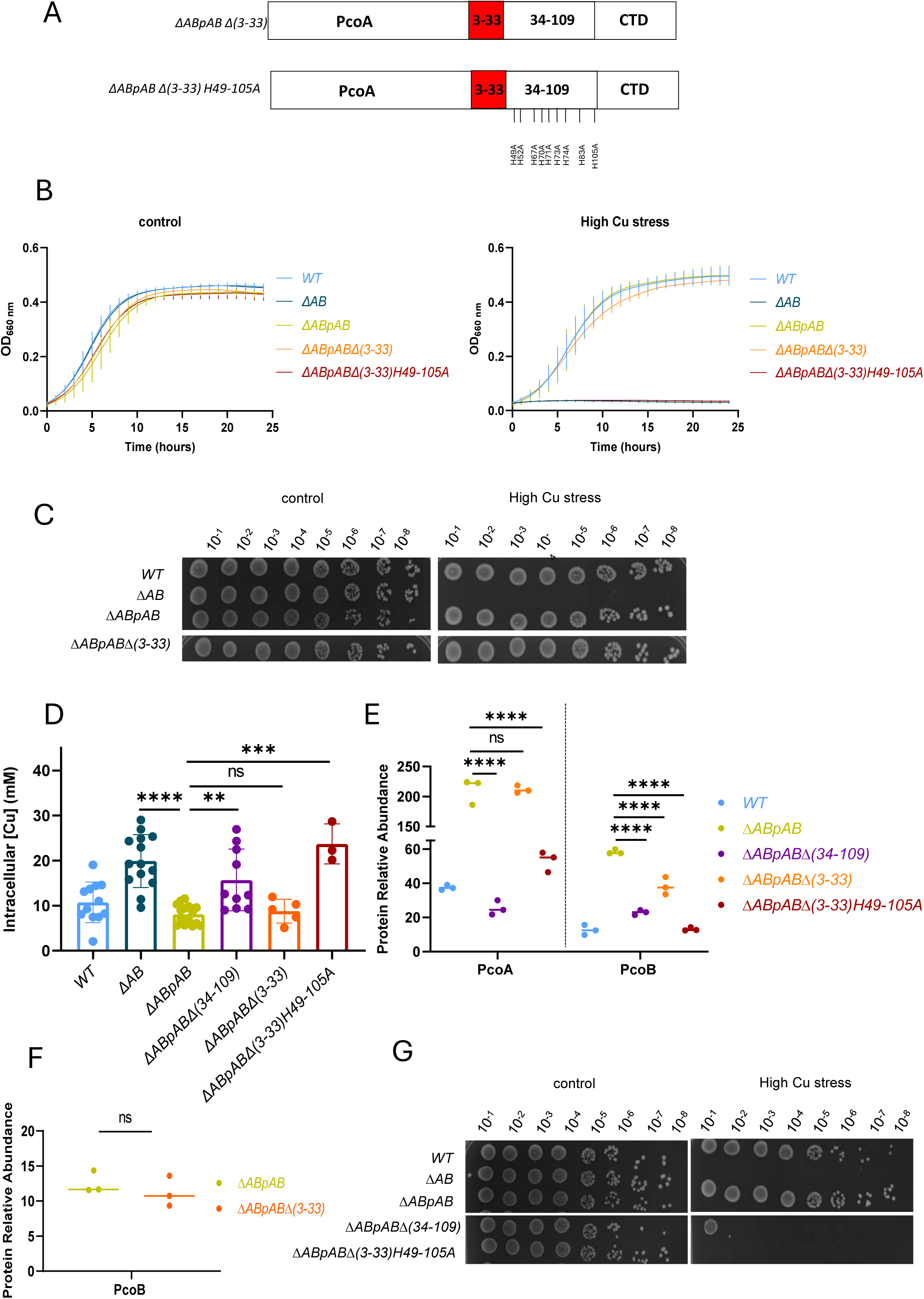
PcoB harbors an unconventional signal peptide. **A.** schematic representation of the *ΔABpABΔ(3-33) and ΔABpABΔ(3-33)H49-105A* mutants. **B.** Growth profiles at an absorbance of 660 nm of WT*, ΔAB, ΔABpAB, ΔABpABΔ(3-33)* and *ΔABpABΔ(3-33)H49-105A* strains grown in PYE medium (left) and in PYE medium supplemented with CuSO_4_ (right). **C.** Viability assay on PYE plates of WT*, ΔAB, ΔABpAB and ΔABpABΔ(3-33)* strains, grown in control and high CuSO_4_ stress conditions. **D**. Average Cu concentration in each bacterial cell in the WT*, ΔAB, ΔABpAB, ΔABpABΔ(34-109), ΔABpABΔ(3-33)*, and *ΔABpABΔ(3-33)H49-105A* strains exposed to 175 µM CuSO_4_ for 5 min. Mean ± SD, at least three biological replicates. **E.** Normalized spectrum counts of PcoA and PcoB peptides in the WT*, ΔABpAB, ΔABpABΔ(34-109), ΔABpABΔ(3-33), and ΔABpABΔ(3-33)H49-105A* strains grown in PYE medium, measured by LC-MS. Individual values and means represented. *p* values were calculated using ANOVA combined with Dunnett’s multiple comparison test (\**p* < 0.05, ** *p* < 0.01, *** *p* < 0.001 and \*\*\*\**p* < 0.0001). **F.** PcoB abundance measured by LC/MS in OM extracts of the *ΔABpAB and ΔABpAB Δ(3-33)* strains grown in PYE medium. *p* values were calculated using an Unpaired t test; \**p* < 0.05. **G.** Viability assay on PYE plates of *WT, ΔAB, ΔABpAB, ΔABpABΔ(34-109) and ΔABpABΔ(3-33)H49-105A* strains, grown in control and high CuSO_4_ stress conditions.

To address the role of the eight His residues located within the PcoB NTD_3-33_, we generated the *ΔABpABΔ(3-33)H49-105A* mutant, which combines the 9 His to Ala point mutations of the PcoB NTD_34-109_ to the loss of the 8 His located in the PcoB NTD_3-33_ **(Fig. 6A**). This mutant phenocopies the *ΔAB* mutant in term of Cu sensitivity in liquid medium and on plate **(Fig. 6B and G**) and Cu content **(Fig. 6D**). The *ΔABpABΔ(3-33)H49-105A* mutant also displayed a significant decrease of PcoB level compared to the *ΔABpABΔ(34-109)* and the *ΔABpAB Δ(3-33)* **(Fig. 6E**). However, the PcoA level is significantly different between the *ΔABpABΔ(34-109)* mutant and the *ΔABpABΔ(3-33)* mutant, suggesting that PcoA stabilization by PcoB does not entirely rely on the His residues present in PcoB NTD.

Altogether, our data highlight the role of the PcoB IDR in the stability and function of PcoA and PcoB and in turn in Cu resistance. They also reveal that PcoB IDR is quite tolerant to sequence and size change, if a minimal set of histidine residues is present.

## Discussion

Copper (Cu) efflux is a central strategy used by organisms to resist Cu toxicity. In the Gram-negative aquatic alphaproteobacterium *C. vibroides*, a chromosomally encoded operon supports Cu homeostasis via PcoA and PcoB, responsible for Cu oxidation and efflux, respectively (17). While the role of PcoA is established, the mechanism of PcoB-mediated Cu efflux remains poorly understood.

The predicted 3D structure of PcoB reveals a 12-stranded antiparallel β-barrel preceded by a highly flexible, intrinsically disordered NTD comprising 109 residues (18). Intrinsically disordered regions (IDRs) are widespread across all domains of life and are crucial for various cellular functions, including signaling, protein interactions, and transcriptional regulation (23).

### Functional Role of the PcoB NTD

Our findings demonstrate that a *ΔABpABΔ(34-109)* mutant, lacking most of PcoB NTD, displays an increased Cu sensitivity and intracellular Cu accumulation, highlighting the NTD critical role in Cu efflux.

The His- and Met-rich motifs 29HAHH32, 49HAGH52, and 67HAGHHMHH74—within residues 34–109 resemble the amino-terminal Cu(II)- and Nickel Ni(II)-binding (ATCUN) motif NH₂–X–X–H, typically found in proteins that bind Cu and Ni (24). Accordingly, site-directed mutagenesis of the 9 His or of the 4 Met in this region disrupts Cu efflux. Furthermore, swapping or cropping the 34–109 region with the corresponding region from *E. coli* PcoB—containing a minimal set of His and Met residues—retains functionality, suggesting these residues are essential.

A comparable system is seen in the human Cu(I) ions transporter hCtr1, where His- and Met-rich motifs in the disordered NTD aid in Cu acquisition through an energy-independent diffusion process (25).

In Gram-negative bacteria, metal efflux typically relies on tripartite HME-RND complexes energized by the proton motive force. However, as an outer membrane (OM) protein lacking known energy sources such as ATP, GTP, or ion gradients, the energy-independent function of PcoB raises questions. One plausible mechanism is a diffusion-based, entropy-driven process similar to the secretion of the curli-forming CsgA by the CsgG/E/F system in *E. coli* (26), wherein the dynamic PcoB NTD would form a transient pocket for Cu trapping and release via Brownian motion.

IDPs often undergo disorder-to-order transitions upon interaction with binding partners (23). In hCtr1, Cu(I) binding induces a transition from disorder to a β-conformation, while interaction with lipids promotes α-helical structure formation (27, 28). This suggests that PcoB’s disordered NTD maintains its flexibility through its His residues and adopts a more structured but transient, state upon Cu binding, facilitating transport.

### NTD Contributions to PcoB and PcoA Stability

PcoB protein levels were significantly reduced in the *ΔABpABΔ(34-109)* mutant, and even more so in the *ΔABpABΔ(3–33)H49–105A* variant lacking all His in the NTD, suggesting that a minimal number of His is critical for PcoB stability. This effect is not due to altered *pcoB* transcription or PcoB half-life.

The four-base overlap between *pcoA* and *pcoB* ORF supports a model of translational coupling. The present study shows that PcoA stability is also compromised in the absence of PcoB NTD, indicating a functional interplay between PcoA and PcoB. This may involve direct interaction, with the disordered NTD serving as a docking site for PcoA, potentially facilitating Cu(II) transfer from PcoA to PcoB, while stabilizing PcoA.

IDPs often form transient yet specific interactions with globular proteins, involving disorder-to-order transitions within 10–50 amino acid segments (29–31). This mechanism may underline the interaction between PcoB and PcoA.

### Unconventional Signal Peptide in Caulobacter PcoB

As an OM protein, PcoB would be expected to possess a cleavable signal peptide (SP) for translocation via the Sec pathway (32). However, our computational and experimental analyses indicate that *C. vibroides* PcoB lacks a classical SP.

This aligns with the phenomenon of “non-classical secretion,” first described in eukaryotes for proteins such as interleukin-1β and thioredoxin (33, 34), and later observed in bacteria, including secretion of GlnA and SodA without Sec or Tat SPs (35, 36).

In *E. coli*, the HybC subunit is co-transported with its Tat-SP-carrying partner HybO via a “piggyback” mechanism (37). Given the potential interaction between PcoA and PcoB, one could hypothesize a similar mechanism where PcoB might undergo translocation through a “hitchhiking” system, potentially coupled to PcoA through the Tat system. However, the presence of PcoB in the periplasmic/OM fraction of the *ΔABpB* mutant lacking PcoA suggests that PcoB translocation occurs independently.

In conclusion, our study reveals an unprecedented role of an IDR in Cu efflux and protein stabilization, while also challenging the traditional paradigm of signal peptide-dependent OM translocation in Gram-negative bacteria.

## Material and Methods

### Strains and plasmids

The *C. vibroides NA1000 (*WT) strain was grown at 30°C, under moderate shaking, in Peptone Yeast Extract (PYE) (Poindexter, 1981) medium, supplemented with 5 μg/ml kanamycin, 15 μg/ml nalidixic acid, 100 μg/ml chloramphenicol and/or CuSO_4_.5H_2_O when required. Exponentially growth cultures were used for all experiments. Plasmids were mobilized from a *DH10B E. coli* strain into *C. vibroides* by triparental mating (Glazebrook & Walker, 1991). Strains and plasmids are listed in Table S1. The strategies and the primers for their construction are available upon request

### Growth curves

Bacterial cultures in the exponential growth phase (OD_660_ of 0.4 – 0.6) were diluted in PYE medium to a final OD_660_ of 0.05 and inoculated in 96-well plates with appropriate CuSO_4_ concentrations when required. Bacteria were then grown for 24 h at 30°C under continuous shaking in an Epoch 2 Microplate Spectrophotometer from BioTek and OD_660_ was measured every 10 min.

### Viability assay

Bacterial cultures in the exponential growth phase (OD_660_ of 0.4 – 0.6) were diluted in PYE medium to a final OD_660_ of 0.1. Ten-fold serial dilutions up to 10^-8^ (in PYE) were prepared in 96-well plates and drops of 5 μL of each dilution were spotted on PYE and PYE Cu plates using an automatic multichannel. Plates were incubated for 48 h at 30°C and pictures were taken with and pictures were taken with the Amersham Imager 600 (GE Healthcare LifeSciences).

### ICP-OES

15 mL cultures of *C. vibroides* cells, were grown up to exponential phase (OD_660_ = 0.5). A 5-minute treatment with 175 μM final CuSO_4_ concentration was applied, and the cultures were then centrifuged at 8500 rpm for 10 min at 4°C (trash supernatant). The cells were then fixed for 20 min on ice in 2% paraformaldehyde and then washed three times with an ice-cold wash buffer (10 mM Tris-HCl pH 6.8, 100 μM EDTA). For a total fraction, where the cytoplasm and periplasm are not separated, the pellet is resuspended in 2 ml of milliQ H_2_O and then lysed with the cell disrupter (Cell Disruption System, One-shot Model, Constant) at 2.48 kPa. A final centrifugation at 10,000 rpm is performed for 10 min and 1.6 mL of the supernatant is mixed with 1 mL HNO_3_ 5% and 2.4 mL milliQ H_2_O for a final volume of 5 mL and a final concentration of 1M HNO_3_. Samples were finally analyzed by ICP-OES with Optima 8000 ICP-OES (PerkinElmer) and cellular metal concentrations were calculated using the following formula as in (17):

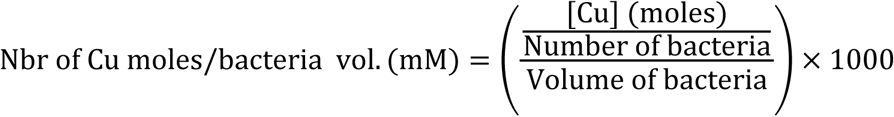

With

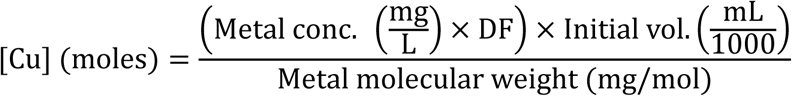

### Mass Spectrometry

#### periplasmic/OM sample preparation

15 mL cultures of *C. vibroides* cells, were grown up to exponential phase (OD_660_ = 0.4-0.6). The cultures were then centrifuged at 8,000 rpm for 10 min at 4°C (trash supernatant) and washed three times with an ice-cold wash buffer (10 mM Tris-HCl pH 6.8, 100 μM EDTA). The pellets were then incubated in 2ml, V/v of Zwittergent 0.25% and Zwittergent buffer (0.2 M Tris-HCl pH 7.6) for 10 min at room temperature and centrifuged for 15 min at 19,000 x g. The supernatant, containing the periplasmic/OM components, was separated from the cytoplasmic pellet and collected as samples for LC/MS analysis.

### OM sample preparation

The OM fraction of *C. vibroides* was harvested by ultracentrifugation after Sodium Lauryl Sarconisate (SLS) and Sodium Carbonate treatment as described in (38). 400 ml of cell culture were grown in PYE at 30°C to an O.D.660 of 0.4. Bacteria were centrifuged for 10 min at 4000 x g at 4°C and washed 3 times in 50 ml of 50 mM (pH 8) Ammonium bicarbonate (AmBic). Cells were finally resuspended in 5 ml AmBic and sonicated on ice (20 rounds of 5 seconds sonication at maximum intensity). The lysate was centrifuged for 20 min at 12,000 x g at 4°C and the pellet containing the unbroken cells and debris was discarded. The supernatant was then ultracentrifuged for 40 min at 100,000 x g to dissociate the cytoplasmic fraction (supernatant) from the total membrane fraction (pellet). The pellet was then resuspended in 1 ml of 1% Sodium Lauryl Sarconisate incubated 30 min at RT and centrifuged for another 40 min at 100,000 x g and 4°C. The subsequent supernatant contains the solubilized inner membrane (SIM) and was discarded. The pellet containing the OM fraction was washed in 1 ml of 2.5 M NaBr and incubated for 30 min on ice. It was then ultracentrifuged for 40 min at 100,000 x g at 4°C. The supernatant was discarded, and the pellet was incubated for 1 h in 1 ml of 100 mM Na_2_CO_3_ for further enrichment of the OM fraction, before being spun again for 40 min at 100,000 x g at 4°C. The OM fractions were collected as samples for LC/MS analysis.

### LC/MS

The samples were treated using the optimized filter-aided sample preparation protocol (39) Briefly, the samples were loaded onto Millipore Microcon 30 MRCFOR030 Ultracel PL-30filters that have been rinsed and washed beforehand with 1% formic acid (FA) and 8 M urea buffer (8 M urea in 0.1 M Tris buffer at pH 8.5), respectively. The proteins on the filter were then exposed to a reducing agent (dithiothreitol) and then alkylated with iodoacetamide. The proteins were then finally digested overnight with trypsin or trypsin/Glu-C (trypsin in 1/50 in ABC buffer; Glu-C in phosphate buffer 50 mM pH 7.4). The final step of digestion is to transfer proteins in 20 μL of 2% acetonitrile (ACN) and 0.1% FA in an injection vial for reverse phase chromatography. The digest was analyzed using nano-LC–ESI–MS/MS timsTOF Pro (Bruker, Billerica, MA, USA) coupled with an UHPLC nanoElute (Bruker). Peptides were separated on a 75 μm ID, 25 cm C18 column with integrated Captive Spray insert (Aurora, Ion Opticks, Melbourne) at a flow rate of 400 nL/min at 50°C. LC mobile phase A was water with 0.1% formic acid(v/v), and B was ACN with formic acid 0.1% (v/v). Samples were loaded directly on the analytical column at a constant pressure of 800 bars. The digest (1 μL) was injected, and the organic content of the mobile phase was increased linearly from 2% B to 15% in 22 min, 15% B to 35% in 38 min, and 35% B to 85% in 3 min. Data acquisition on the tims TOF Pro was performed using Hystar 5.1 and tims Control 2.0. tims TOF Pro data were acquired using 100 ms TIMS accumulation time and mobility (1/K0) range from 0.6 to1.6 Vs/cm². Mass-spectrometric analysis was carried out using the parallel accumulation serial fragmentation (PASEF) acquisition method (40). One MS spectrum was followed by 10 PASEF MSMS spectra per total cycle of 1.1 s. All MS/MS samples were analyzed using Mascot (Matrix Science, London, UK; version 2.8.1).

Mascot was set up to search the *C. vibroides* NA1000_190306 database from UniRef 100 and Contaminants_20190304 database assuming the digestion enzyme trypsin/GluC. Mascot was searched with a fragment ion mass tolerance of 0.050 Da and apparent ion tolerance of 15 PPM. Carbamidomethyl of cysteine was specified in Mascot as a fixed modification. Oxidation of methionine and acetyl of the n-terminus was specified in Mascot as variable modifications. Scaffold (version Scaffold_5.1.1, Proteome Software, Inc., Portland, OR) was used to validate MS/MS-based peptide and protein identifications. Peptide identifications were accepted if they could be established at greater than 97.0% probability to achieve an FDR less than 1.0% by the Percolator posterior error probability calculation (41). Protein identifications were accepted if they could be established at greater than 50.0% probability to achieve an FDR less than 1.0% and if they contain at least one identified peptide. Protein probabilities were assigned by the Protein Prophet algorithm (42). Proteins that contained similar peptides and could not be differentiated based on MS/MS analysis alone were grouped to satisfy the principles of parsimony. Proteins sharing significant peptide evidence were grouped into clusters.

Given that we are working with different lengths of the PcoB protein, only peptides corresponding to the CTD (residues 111–302) were included in the quantification analysis. Peptides from the NTD were excluded from the Scaffold file to ensure consistent comparison of peptide counts.

### Western blotting

Exponentially growing cells were collected and re-suspended in an SDS-PAGE loading buffer, with the pellet volumes adjusted to achieve an OD_660_ of 1, ensuring protein content normalization. The protein samples were then boiled and separated on 12% sodium dodecyl sulfate-polyacrylamide gels before being electro-transferred to a nitrocellulose membrane. Subsequently, the nitrocellulose membranes were probed with polyclonal rabbit anti-PcoA (1:5000) and anti-PcoB (1:1000). A polyclonal goat anti-rabbit immunoglobulins/HRP secondary antibody at a dilution of 1:10,000 (DAKO) was employed.

Cy5 fluorescence was used as a loading control by adding 1 µl of Amersham Cy5 dye to the protein samples, followed by incubation for 30 min at room temperature and heating for 5 min at 95 °C. The Cy5 fluorescence was detected using the Amersham Imager 600.

### RT-qPCR

Bacteria were grown in PYE up to OD660nm = 0.4 before incubation at 30°C under agitation. Bacteria were recovered by centrifugation, and pellets were flash-frozen until resuspension in 40 μL of a 20 mg/mL proteinase K solution (Avantor, Radnor, PA, USA) with 1 μL of undiluted Ready-Lyse Lysozyme Solution (Lucigen, Middlesex, UK), and lysis was allowed to proceed for 10 min in a shaking incubator at 37°C and 600 rpm. Total RNA was retrieved from the cell suspensions using TriPure isolation reagent and procedure as described by the manufacturer (Roche, Mannheim, Germany). RNA (2 μg) isolated from *C. vibroides* was incubated with DNase I (Thermo Scientific, Merelbeke, Belgium) for 30 min at 37°C. DNase I was then inactivated with 50 mM EDTA for 10 min at 65°C. Subsequently, RNA was subjected to reverse transcription using MultiScribe Reverse Transcriptase (Applied Biosystems, Foster City, CA, USA) with random primers (as described by the manufacturer). A total of 300 ng of cDNA was mixed with Takyon No Rox SYBR MasterMix dTTP Blue (Eurogentec, Seraing, Belgium) and the appropriate primer sets were used for qPCR in LightCycler96 (Roche, Basel, Switzerland). Forty-five PCR cycles were performed (95°C for 10 s, 60°C for 10 s, and 72°C for 10 s). Primer specificity was checked by melting curve analysis. Relative gene expression levels between different samples were calculated with the 2−ΔΔCt method using the rpoD gene as a reference. Four biological replicates were analyzed for each sample.

The sequences of the primers are available on request.

### Protein half-life

Exponentially growing cells (OD_660_ = 0.4-0.6) were treated with 100 μg/ml chloramphenicol to arrest protein synthesis. Samples were collected at 1 h, 2 h and 4 h after the antibiotic treatment. Pellet volumes were adjusted to achieve an OD_660_ of 1 in 2 ml of milliQ H_2_O, ensuring protein content normalization at each time point. Total fraction of each time point was generated using the cell disrupter (Cell Disruption System, One-shot Model, Constant) at 2.48 kPa. A final centrifugation at 10,000 rpm is performed for 10 min and 1.5 mL of the supernatant was collected. Samples were analyzed with LC/MS.

### Statistical analysis

Statistical analyses were performed when required. The data were analyzed with ANOVA one-way combined with Dunnett’s multiple comparisons test. A *p*-value below 0.05, 0.01, 0.001 and 0.0001 are represented by *, **, ***, and **** respectively. A T-test was performed when needed.

A linear regression statistical test was performed for the Protein half-life experiment.

### Sequence-based bioinformatic predictions

A multiple sequence alignment of PcoB protein from *Caulobacter vibrioides* (Uniprot ID: A0A0H3C699) and 13 homologs from other bacterial species — *Hyphomonas* sp. (Uniprot ID:A0A922YF47), *Sphingobium cupriresistens* (Uniprot ID: A0A0J7XK94), *Rhizobium* sp. (UniParcID: UPI0022C4F4F1), *Methylobacterium aquaticum* (Uniprot ID: A0A0J6SI10), *Nitrobacter* sp. (NCBI ID: WP_319798384), *Bordetella bronchiseptica* (UniParcID: UPI00045A6DA4), *Pseudomonas aeruginosa* (UniParcID: UPI00377290C9), *Enterobacter cloacae* (NCBI ID: WP_185812793), *Salmonella enterica* (UniParcID: UPI001DC5DD2D ), *Klebsiella pneumoniae* (NCBI ID: WP_183406058), *Klebsiella oxytoca* (Uniprot ID: A0A6C0L247), *Acinetobacter baumannii* (Uniprot ID : A0A142G3V9) and *Escherichia coli* (Uniprot ID: Q47453) — was first performed using the online MAFFT tool (https://www.ebi.ac.uk/jdispatcher/msa). The results were visualized using Jalview.

Intrinsic disorder predictions were carried out using the Rapid Intrinsic Disorder Analysis Online (RIDAO) platform (43), which integrates seven well-established per-residue disorder predictors: PONDR VLXT, VL3, VSL2B, PONDR FIT, IUPred-Short, IUPred-Long, and ANCHOR2. The mean disorder profile (σ(MDP)), calculated as the average of the scores from the seven core algorithms mentioned above, was used to generate the disorder plots. Residues with a mean score above 0.5 were considered disordered, whereas those with a score below that threshold were classified as structured.

## Acknowledgements

We thank Valérie Charles and Carmela Aprile (CMI laboratory, NISM, UNamur) for the ICP measurements. We acknowledge Laurelenn Hennaux and the URBM members for fruitful discussions.

## Author contributions

A. K. and J.-Y. M. conceptualization; A. K. methodology; A. K. validation; A. K., M. O., H. B., M. D., P. R., C. M. and J.-Y. M. formal analysis; A. K., M. O., H. B., M. D., investigation; A. K. writing–original draft; A. K. visualization; J.-Y. M. writing–review & editing; J.-Y. M. supervision; J-.Y. M. funding acquisition.

## Figure legends

**Figure S1. Prediction of a signal peptide in PcoB. A.** SignalP5 analysis of PcoB protein sequence. **B.** Manual analysis of PcoB protein sequence. **C.** Sequenced peptide of the Trypsin-digested PcoB from a periplasm/OM fractionate.

**Figure S2. Concentration-dependent Cu sensitivity of the *ΔABpAB Δ(34-109)* mutant. A.** Growth profiles at an absorbance of 660 nm of the WT*, ΔAB, ΔABpAB, ΔABpABΔ(34-109) and BΔ(34-109)* strains grown in PYE medium (top left) and in PYE medium supplemented with increasing CuSO_4_ concentrations. **B.** Viability assay on PYE plates of WT, *ΔAB, ΔABpAB, BΔ(34-109) and ΔABpABΔ(34-109)* strains under high (100 µM) CuSO_4_ stress conditions.

**Figure S3. PcoA and PcoB interdependence.** Normalized spectrum counts of PcoA and PcoB peptides in the WT*, ΔABpAB, ΔABpA and ΔABpB* strains grown in PYE medium, measured by LC-MS. Individual values and means represented. *p* values were calculated using ANOVA combined with Dunnett’s multiple comparison test (\**p* < 0.05, ** *p* < 0.01, *** *p* < 0.001 and \*\*\*\**p* < 0.0001)

**Figure S4. PcoB half-life.** PcoB abundance measured by LC/MS in total cell extracts obtained 0 h, 1 h, 2 h and 4 h after chloramphenicol treatment of the *ΔABpAB and ΔABpABΔ(34-109)* strains grown in PYE medium. *p* values were calculated using a linear regression statistical test; \**p* < 0.05.

**Figure S5. The predicted signal peptide is not cleaved.** Sequenced peptide of the trypsin/ GluC-digested PcoB from a periplasm/OM fractionate

**Table S1:** Strains and plasmids.

| **Strains** | **Strains Relevant genotype or description** | **References** |
| --- | --- | --- |
| $\Delta AB$ | Knockout strain for *pcoAB* genes | (17) |
| $\Delta ABpA$ | Knockout strain for *pcoAB* genes carrying a copy of *pcoA* on the *pMR10* under the control of the *lac* promoter; $Kan^R$ | (17) |
| $\Delta ABpB$ | Knockout strain for *pcoAB* genes carrying a copy of *pcoB* on the *pMR10* under the control of the *lac* promoter; $Kan^R$ | (17) |
| $\Delta ABpAB$ | Knockout strain for *pcoAB* genes carrying a copy of *pcoAB* on the *pMR10* under the control of the *lac* promoter; $Kan^R$, | (17) |
| $\Delta ABpAB \Delta(34-109)$ | $\Delta ABpAB$, the NTD of *pcoB* is deleted from aa 34 to aa 109 | This study |
| $B \Delta(34-109)$ | The *pcoB* gene on the chromosome is deleted for the NTD from aa 34 to aa 109 | |
| $\Delta ABpAB \Delta(34-59)$ | $\Delta ABpAB$ combined with deletion of the NTD of *pcoB* from aa 34 to aa 59 | This study |
| $\Delta ABpAB \Delta(34-85)$ | $\Delta ABpAB$ combined with deletion of the NTD of *pcoB* from aa 34 to aa 85 | This study |
| $\Delta ABpAB N_{Ec}$ | $\Delta ABpAB$ combined with replacement of the NTD of *pcoB* of *C.c* from aa 34 to aa 109 with the NTD of *pcoB* of *E.c* from aa 25 to aa 90 | This study |
| $\Delta ABpAB H49-105A$ | $\Delta ABpAB$ combined with point mutations of the His 49,52,67,70,71,73,74,83,105 of the NTD of *pcoB* into Ala | This study |
| $\Delta ABpAB M54-102A$ | $\Delta ABpAB$ combined with point mutations of the Met 54,72,94, 102 of the NTD of *pcoB* into Ala | This study |
| $\Delta ABpAB \Delta(3-33)$ | $\Delta ABpAB$ combined with the deletion of aa 3 to aa 33 from the NTD of *pcoB* | This study |
| $\Delta ABpAB \Delta(3-33) H49-105A$ | $\Delta ABpAB$ combined with the deletion of aa 3 to aa 33 from the NTD of *pcoB* and point mutations of the His 49,52,67,70,71,73,74,83,105 into Ala | This study |

